# Modulating Inflammation in Post-Traumatic Osteoarthritis using iPSC-derived Anti-inflammatory Macrophages

**DOI:** 10.64898/2026.05.18.726078

**Authors:** Nadia Mahmoudi, Lea Zila, Julia Sheyn, Namdev More, Melissa Chavez, Dean Roell, Rachel Lev-Gur, Anoushka Prasad, Sarah Mohyeddinipour, Mia Orr, Minoo Bastani, Oksana Shelest, Sean Rajaee, Andrew Spitzer, Wafa Tawackoli, Dmitriy Sheyn

**Affiliations:** Orthopaedic Stem Cell Research Laboratory, Cedars-Sinai Medical Center, Los Angeles, CA; Board of Governors Regenerative Medicine Institute, Cedars-Sinai Medical Center, Los Angeles, CA; Department of Biomedical Sciences, Cedars-Sinai Medical Center, Los Angeles, CA; Department of Orthopedics and Sports Medicine, Cedars-Sinai Medical Center, Los Angeles, CA; Department of Surgery, Cedars-Sinai Medical Center, Los Angeles, CA

## Abstract

Post-traumatic osteoarthritis (PTOA) is a common long-term consequence of joint injury and a major cause of chronic pain and disability, yet no disease-modifying therapies are currently available. A central barrier to effective intervention is the persistence of maladaptive synovial inflammation, driven in part by macrophage-mediated signaling that sustains tissue degeneration and pain. Here, we developed a scalable, chemically defined platform to generate human induced pluripotent stem cell (iPSC)–derived anti-inflammatory macrophages (iMac-M2) as an off-the-shelf cell therapy designed to restore joint immune homeostasis after injury. These cells maintained a stable anti-inflammatory phenotype and function under osteoarthritis-relevant inflammatory conditions and suppressed inflammatory and catabolic responses in human joint cell co-culture systems. In a preclinical model of PTOA, intra-articular delivery of iMac-M2 after injury improved functional and structural outcomes while modulating synovial inflammatory and pain-associated transcriptional programs. Treatment was well tolerated, with no evidence of systemic immune activation or ectopic tissue formation. Together, these findings support iPSC-derived macrophage therapy as a clinically translatable immunomodulatory strategy to interrupt early inflammatory drivers of PTOA and preserve joint health following injury.

## Introduction

Post-traumatic osteoarthritis (PTOA) is a common and debilitating outcome of joint injuries, leading to chronic pain, functional disability, and substantial economic burden due to its high prevalence and costly management.^1^ Approximately 12% of all osteoarthritis cases are attributed to previous joint trauma,^1^ including fractures,^2^ ligament tears such as anterior cruciate ligament (ACL) ruptures^2^ and meniscal damage.^3^ Joint trauma initiates a cascade of biomechanical disruption and inflammatory signaling that progressively compromises joint integrity, frequently culminating in PTOA. Notably, up to 50% of joint injuries progress to PTOA, typically manifesting within 7-10 years after the initial trauma.^4^

Unlike primary osteoarthritis, which predominantly affects older individuals, PTOA impacts patients across all age groups, with young adults and athletes being particularly vulnerable due to their high exposure to joint injuries.^5^ Recent studies underscore the alarming rise in PTOA prevalence, with projections showing it could affect as many as 40.6% of individuals by 2030, up from 21.1% in 2022.^6^ These trends highlight the clinical importance of early disease-modifying interventions aimed at altering disease trajectory rather than managing end-stage degeneration.^7^

PTOA is now recognized as a whole-joint disease, involving coordinated pathological changes in cartilage, subchondral bone, synovium, and periarticular tissues.^7^ Following joint injury, an acute inflammatory response is triggered, characterized by elevated levels of pro-inflammatory cytokines such as IL-1β, IL-6, IL-8, and TNF-α in the synovial fluid.^8^ While this response is initially required for tissue repair, its persistence promotes matrix metalloproteinase (MMPs) activation and cartilage degradation.^9^

Importantly, a subset of patients exhibits a dysregulated inflammatory response following ACL injury, marked by sustained elevation of pro-inflammatory mediators and cartilage degradation markers.^10^ Such unresolved inflammation is a major risk factor for PTOA development, suggesting that early immunomodulatory intervention may be critical for disease prevention.^8^

Although surgical techniques, such as ligament reconstruction, can restore joint stability, their impact on PTOA prevention remains limited.^11,12^ This limitation underscores a critical gap between biomechanical correction and long-term biological joint protection, indicating that mechanical restoration alone is insufficient to halt PTOA progression. Increasing evidence highlights the central role of pro-inflammatory cytokines and MMPs in sustaining cartilage degeneration and tissue damage following joint injury.^13^

Blunt cartilage compression during ACL injuries induces bone bruises and osteochondral microtrauma, associated with inflammatory cytokine dysregulation and early cartilage injury. These events initiate prolonged inflammatory cascades, with pathways such as IL-1 signaling playing a key role in PTOA onset and progression.^14^ Similarly, displaced intra-articular fractures exacerbate disease development by activating inflammatory pathways and promoting chondrocyte death, accelerating cartilage destruction.^15^ Mechanical loading further amplifies these effects by altering matrix metabolism and enhancing the production of inflammatory mediators, including nitric oxide and prostaglandin E2.^16^ While surgical interventions such as ACL repair restore partial mechanical stability, they frequently fail to resolve these inflammatory and catabolic processes, thereby limiting their ability to prevent PTOA progression.^17^ Bone bruises and osteochondral damage, commonly observed after acute joint trauma, illustrate the tight interplay between biomechanical injury and inflammation-driven cartilage degeneration, although their precise contribution to long-term functional outcomes remains under investigation.^18,19^

The immune system plays a pivotal role in PTOA pathogenesis, with macrophages emerging as central regulators of post-injury inflammation.^20^ Synovial macrophages, which normally contribute to joint homeostasis through debris clearance and tissue repair, become aberrantly activated following injury.^21^ This activation leads to an imbalance between pro-inflammatory-like (M1) macrophages and anti-inflammatory, pro-regenerative-like (M2) macrophages.^22^

The dominance of M1 macrophages sustains chronic inflammation, driving synovial fibrosis, cartilage breakdown, subchondral bone damage, and persistent pain.^14,23^ Although cytokine levels often decline over time, certain mediators such as IL-6 and TNF-α can remain elevated for months or years, while anti-inflammatory regulators such as IL-1Ra may remain insufficient.^24,25^ Failure to resolve post-traumatic inflammation therefore represents a key mechanistic driver of PTOA progression.^23,26^

Restoring macrophage polarization balance has thus emerged as a promising therapeutic strategy for PTOA. By enhancing M2 macrophage activity while limiting M1-driven inflammation, it may be possible to promote tissue repair, mitigate pain, and preserve joint structure.^27^ However, current PTOA treatments remain largely surgical and symptomatic, providing limited protection against long-term disease progression.^28^

Induced pluripotent stem cells (iPSCs) provide a scalable and clinically relevant platform for regenerative immunotherapies, enabling the controlled generation of specialized immune cells at scale.^29–31^ iPSC-derived M2 macrophages (iMac-M2) represent a particularly attractive approach, as they secrete anti-inflammatory cytokines (e.g., IL-10, TGF-β), suppress catabolic mediators such as TNF-α and IL-1β, and promote tissue repair through growth factor release.^32,33^ By directly reprogramming the post-injury immune environment, iMac-M2 therapy has the potential to intervene at an early, disease-modifying stage of PTOA. In parallel, in vitro PTOA models using inflamed human chondrocytes and synoviocytes provide valuable platforms for mechanistic studies and therapeutic evaluation ^34^. These systems enable controlled investigation of inflammatory signaling, matrix degradation, and immune–stromal cell interactions relevant to PTOA pathogenesis.

Pain represents a defining and debilitating feature of PTOA, arising from the combined effects of inflammation, structural joint damage, and aberrant innervation.^35,36^ Inflammatory mediators sensitize peripheral nociceptors, while abnormal nerve sprouting, including increased calcitonin gene-related peptide (CGRP)-positive fibers, amplifies pain signaling within the joint.^37,38^

The destabilization of the medial meniscus (DMM) model is a well-established preclinical model that recapitulates key features of human PTOA, including joint instability, progressive cartilage degeneration, and pain-related behaviors.^39,40^ Given the higher prevalence and accelerated progression of PTOA in males, this study employs male rats to reflect clinically relevant disease dynamics.

We hypothesize that early intra-articular administration of iPSC-derived M2 macrophages can prevent or delay PTOA onset by rebalancing post-traumatic inflammation, limiting tissue degeneration, and reducing pain.

Accordingly, the objective of this study is to evaluate the immunomodulatory, structural, and functional effects of iMac-M2 therapy using complementary in vitro PTOA models and an in vivo DMM rat model, thereby assessing its potential as a disease-modifying strategy for PTOA.

## Results

### Differentiation and characterization of anti-inflammatory macrophages (iMac-M2)

Human induced pluripotent stem cells (iPSCs) were successfully differentiated into anti-inflammatory M2 macrophages (iMac-M2) through a stepwise protocol using chemically defined media and small molecules (Fig. 1A-B). The differentiation process involved sequential stages including iMyeloid progenitors, CD14⁺ iMonocytes, CD68⁺ macrophages, and final polarization into M2 macrophages characterized by CD163⁺/CD206⁺ expression. The differentiation protocol was reproducible across multiple independent iPSC lines, yielding consistent cellular populations at each stage of differentiation (Fig. S1).

**Fig 1.**
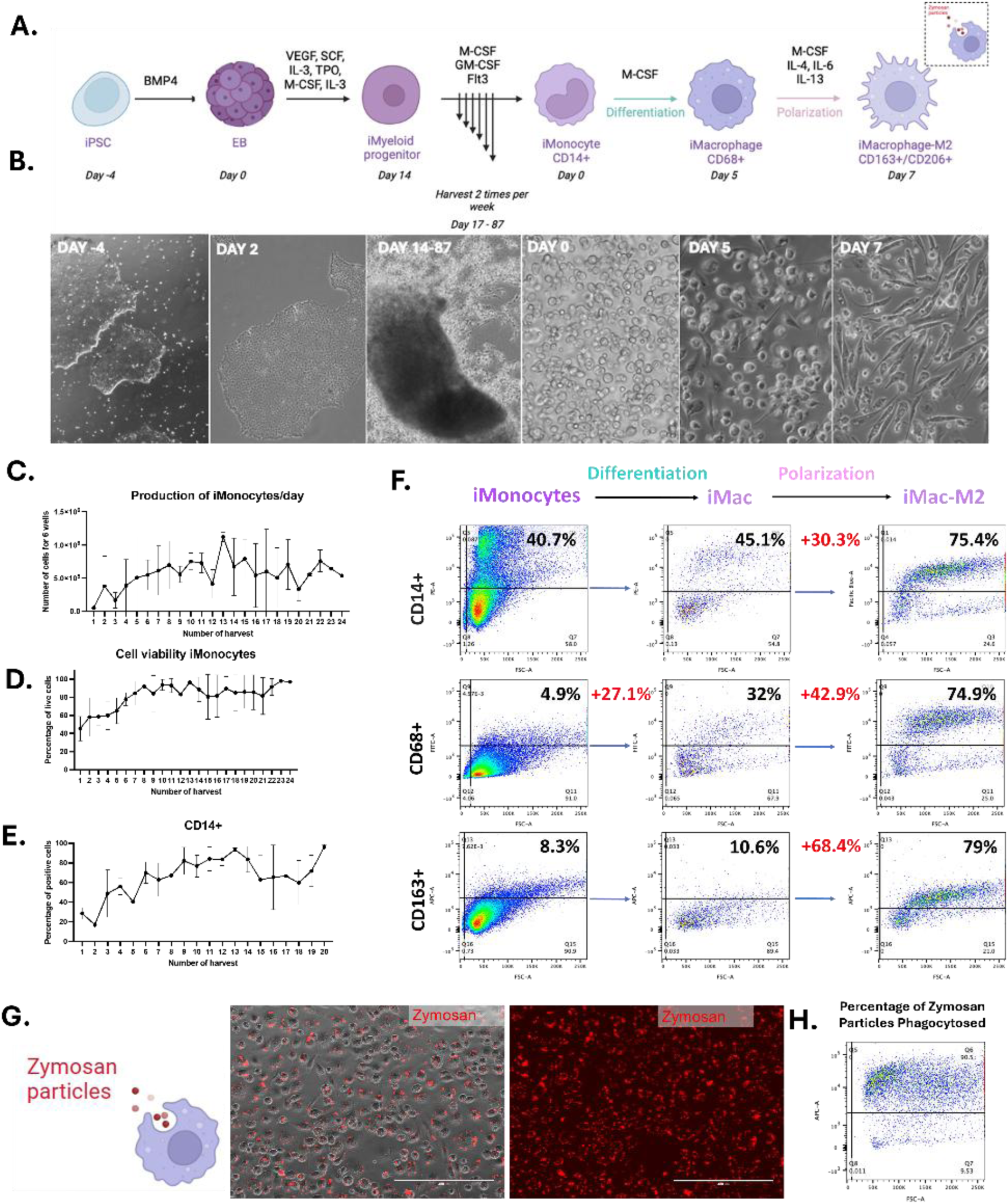
Generation and characterization of anti-inflammatory iPSC-derived macrophages (iMac-M2). **(A)** Schematic overview of the stepwise differentiation protocol used to generate anti-inflammatory macrophages from induced pluripotent stem cells (iPSCs) using chemically defined media and cytokine supplementation. iPSCs first form embryoid bodies (EBs), which give rise to myeloid progenitors and subsequently CD14⁺ iMonocytes. These cells are further differentiated into macrophages (iMac) and polarized toward an anti-inflammatory phenotype (iMac-M2). (B) Representative brightfield images illustrating cellular morphology at successive stages of differentiation, including EB formation, iMonocyte production, macrophage differentiation, and M2 polarization. (C) Quantification of the number of iMonocytes harvested per day during the production phase. (D) Viability of harvested iMonocytes during culture. (E) Flow cytometry analysis confirming the expression of the monocyte marker CD14 in iPSC-derived iMonocytes. (F) Representative flow cytometry (FACS) plots showing the expression of the M2-associated markers CD163 and CD206 following macrophage differentiation and polarization, confirming the generation of iMac-M2 cells. (G) Representative fluorescence microscopy images demonstrating the phagocytic activity of iMac-M2 cells after incubation with Zymosan particles, with corresponding dot plots. (H) Quantification of the percentage of Zymosan particles phagocytosed by iMac-M2 cells. Data in C-E are presented as mean ± SD. N = 2, where N represents the number of independent differentiation experiments performed with iPSC line CS0007iCTR-n5.

Daily harvests of iMonocytes showed stable production with high viability (>90%) over time (Fig. 1C-D). The proportion of CD14⁺ cells progressively increased across successive harvests (Fig. 1E), indicating sustained and efficient generation of iMonocytes.

Following polarization with IL-4, IL-6 and IL-13, the resulting iMac-M2 population exhibited a marked increase in the expression of M2-associated markers, with 79% of cells expressing CD163 (Fig. 1F). Functional characterization of iMac-M2 demonstrated robust phagocytic activity (Fig. 1G), with more than 90% of cells internalizing Zymosan particles (Fig. 1H).

### Distinct Transcriptional profiles of iMac-M2 from iMac

Gene expression analysis of iMac and iMac-M2 markers revealed distinct transcriptional profiles associated with inflammatory and tissue-regulatory states. The heatmap analysis (Fig. 2) showed that iMac-M2 macrophages, compared to unpolarized macrophages (iMac), exhibited increased expression of M2-associated and anti- inflammatory genes, including CD163, CD209, MRC1, PPARG, IL10, CCL13, CCL18, CLEC10A, and ALOX15. Conversely, iMac samples showed higher expression of genes associated with inflammatory activation, including MMP9, CSF1R, and CCR2. Genes associated with macrophage motility and tissue surveillance, including CX3CR1 and FCMR, showed comparable expression levels between iMac and iMac-M2 samples, indicating that polarization was not associated with major transcriptional changes in mobility-related pathways. In addition, genes involved in tissue remodeling and extracellular matrix regulation, such as MMP10 and MMP12, were differentially expressed between iMac and iMac-M2s, consistent with distinct transcriptional programs associated with inflammatory versus tissue-regulatory macrophage states.

**Fig 2.**
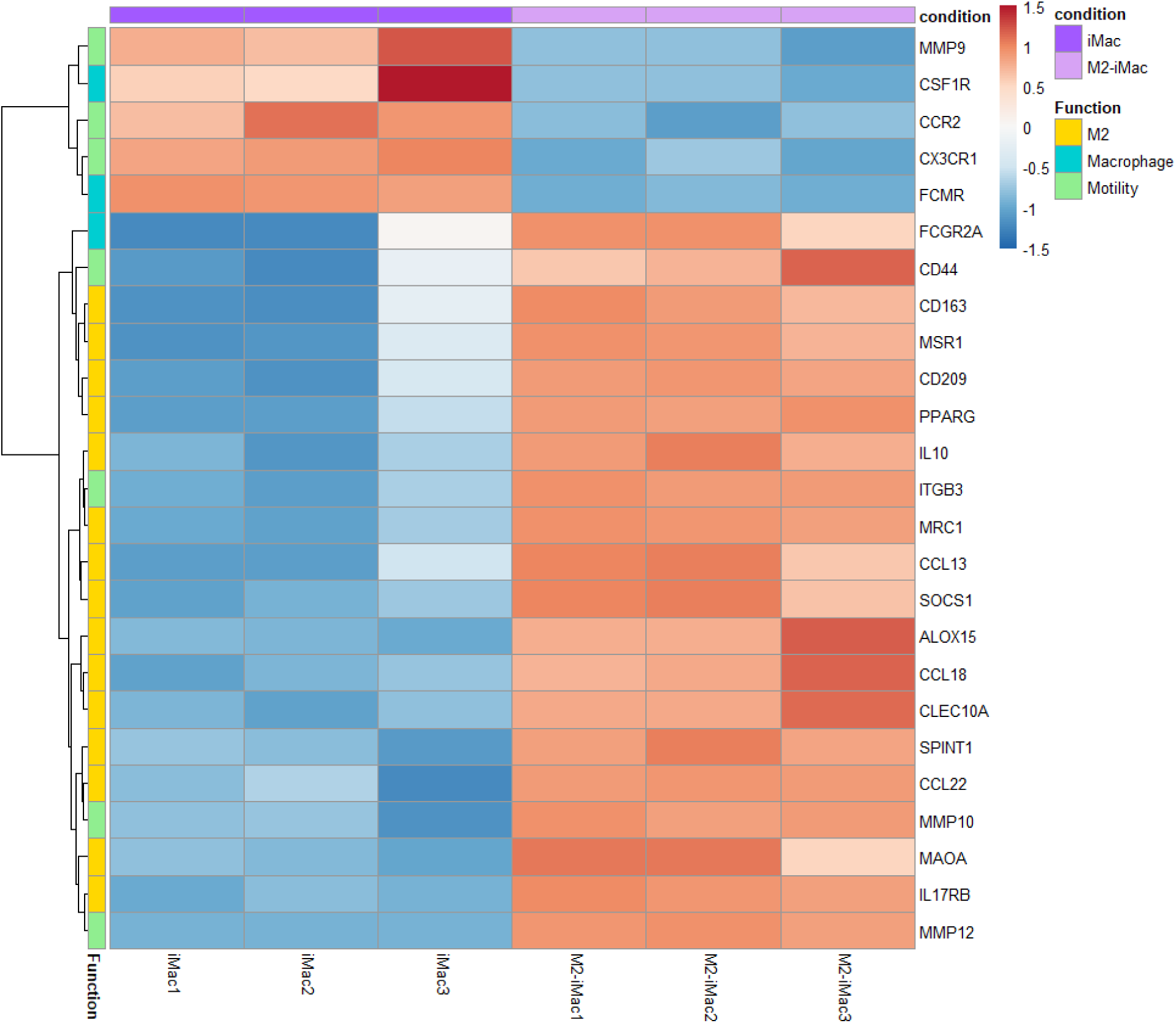
Transcriptomic profiling reveals an M2-associated gene expression signature in iMac-M2 cells. Heatmap showing the expression of macrophage-related genes in iPSC-derived macrophages (iMac) and polarized iMac-M2 cells based on bulk RNA sequencing analysis. (iMac1–3 and M2-iMac1–3). Gene expression values are displayed as normalized and scaled expression levels across samples. Hierarchical clustering highlights distinct transcriptional profiles between iMac and M2-iMac cells. Genes associated with macrophage motility, macrophage identity, and M2 polarization are indicated by functional annotation bars. iMac-M2 cells show increased expression of canonical M2-associated genes, including CD163, MRC1 (CD206), MSR1, IL10, PPARG, CCL18, CCL22, and ALOX15, consistent with an anti-inflammatory macrophage phenotype (n = 3 per condition). N represents the number of separate culture plates derived from the same iPSC line CS0007iCTR-n5.

### Stable iMac-M2 phenotype in culture

To assess the temporal stability of iMac-M2 in culture, flow cytometry analysis was performed at multiple time points post-differentiation, from Day 7 to Day 34. The expression of CD14, CD68, and CD163 was maintained across all analyzed time points, indicating persistence of the macrophage and M2-associated phenotype during prolonged culture. Across the culture period, iMac-M2 consistently displayed high levels of CD68 and CD163, with no marked loss of marker expression observed up to Day 34.

### iMac-M2 maintain phenotype under inflammatory conditions

To assess the phenotypic stability of iMac-M2 under inflammatory conditions, cells were exposed to IL-6, TNF-α, IL- 1β, or LPS, which are inflammatory mediators commonly associated with osteoarthritic environments (Fig. 3A). Phase contrast imaging revealed largely preserved morphology across different inflammatory conditions, with only modest morphological changes observed depending on the stimulus and duration of exposure (Fig. 3B). Flow cytometry analysis showed that iMac-M2 retained expression of the M2-associated surface markers CD163 and CD206 at Days 3, 7, and 14 across all inflammatory conditions, although stimulus-dependent variations were observed (Fig. 3C-D). A partial reduction in CD163 and CD206 expression was detected following prolonged exposure to TNF-α or LPS, but M2 marker expression remained detectable at all time points analyzed (Fig. 3C-D).

**Fig 3.**
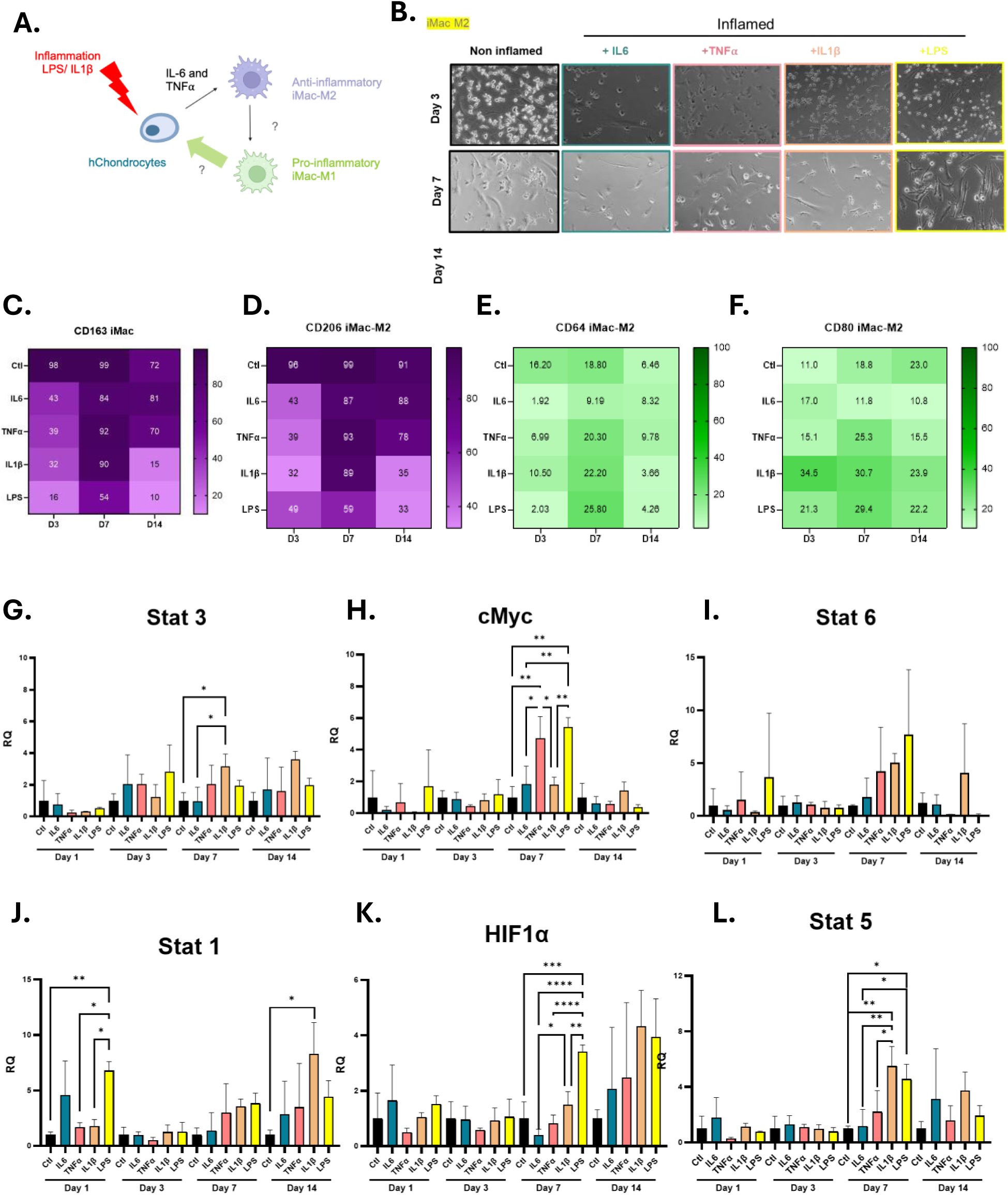
Phenotypic stability of iMac-M2 macrophages under inflammatory conditions. (**A**) Schematic overview of the experimental hypothesis. Inflamed chondrocytes stimulated with IL-1β and LPS adopt an osteoarthritis-like phenotype and release pro-inflammatory cytokines, including IL-6 and TNFα. These inflammatory mediators may influence iPSC-derived M2 macrophages (iMac-M2), potentially promoting a shift toward a pro-inflammatory M1-like phenotype. (**B**) Representative brightfield images showing iMac-M2 morphology under control (non-inflamed) conditions or following exposure to inflammatory stimuli (IL-6, TNFα, IL-1β, or LPS) at day 3, day 7, and day 14. (**C**–**D**) Flow cytometry analysis of M2-associated markers CD163 and CD206 expressed by iMac-M2 at days 3, 7, and 14 after exposure to inflammatory stimuli. (**E**–**F**) Flow cytometry quantification of M1-associated markers CD64 and CD80 under the same conditions and time points. (**G**–**I**) RT–qPCR analysis of M2-associated transcriptional regulators Stat3, cMyc, and Stat6 across inflammatory conditions and time points. (**J**–**L**) RT–qPCR analysis of M1-associated genes Stat1, HIF1α, and Stat5, showing stimulus- and time-dependent transcriptional responses. Data are presented as relative quantification (RQ) compared with control (non-inflamed) iMac-M2, n = 3. Statistical significance is indicated as *P < 0.05, **P < 0.01, ***P < 0.001, ****P < 0.0001. N represents the number of separate culture plates derived from the same iPSC line CS0007iCTR-n5.

In parallel, expression of the M1-associated surface markers CD64 and CD80 remained low overall, with transient increases observed depending on the inflammatory stimulus and time point (Fig. 3E-F). Gene expression analysis further characterized the transcriptional response of iMac-M2 under inflammatory conditions. Expression of M2- associated genes Stat3, cMyc, and Stat6 showed dynamic regulation across stimuli and time points, with sustained expression detected under all inflammatory conditions (Fig. 3G-I). M1-associated genes Stat1, HIF1α, and Stat5 exhibited stimulus- and duration-dependent regulation, with increased expression observed under specific inflammatory conditions (Fig. 3J-L). However, these transcriptional changes did not correspond to a full phenotypic shift, as iMac-M2 cells largely retained expression of M2-associated surface markers. Notably, despite transcriptional modulation of selected M1-associated genes, flow cytometry analyses did not reveal a complete transition of iMac- M2 toward an M1-dominant surface marker profile.

### iMac-M2 reduce inflammation and matrix degradation

To evaluate the effects of iMac-M2 on joint-resident cells in vitro, primary human chondrocytes and fibroblast-like synoviocytes (FLS) isolated from surgical discards of osteoarthritic patients were used (Fig. 4A). Chondrocytes and FLS were exposed to IL-1β and LPS for 24 hours to induce an inflammatory state, followed by direct co-culture with iMac-M2 (Fig. 4A). Acute inflammatory responses were assessed by measuring gene expression of pro-inflammatory cytokines 24 hours after iMac-M2 treatment. To ensure cell type specific gene expression analysis, iMac-M2 were depleted by flow cytometric sorting prior to RNA extraction, enabling selective analysis of chondrocytes and FLS (Fig. S3). The results showed that iMac-M2 significantly reduced the expression of IL-6, IL-1B, and IL-8 in inflamed chondrocytes (Fig. 4B) and FLS (Fig. 4C).

**Fig 4.**
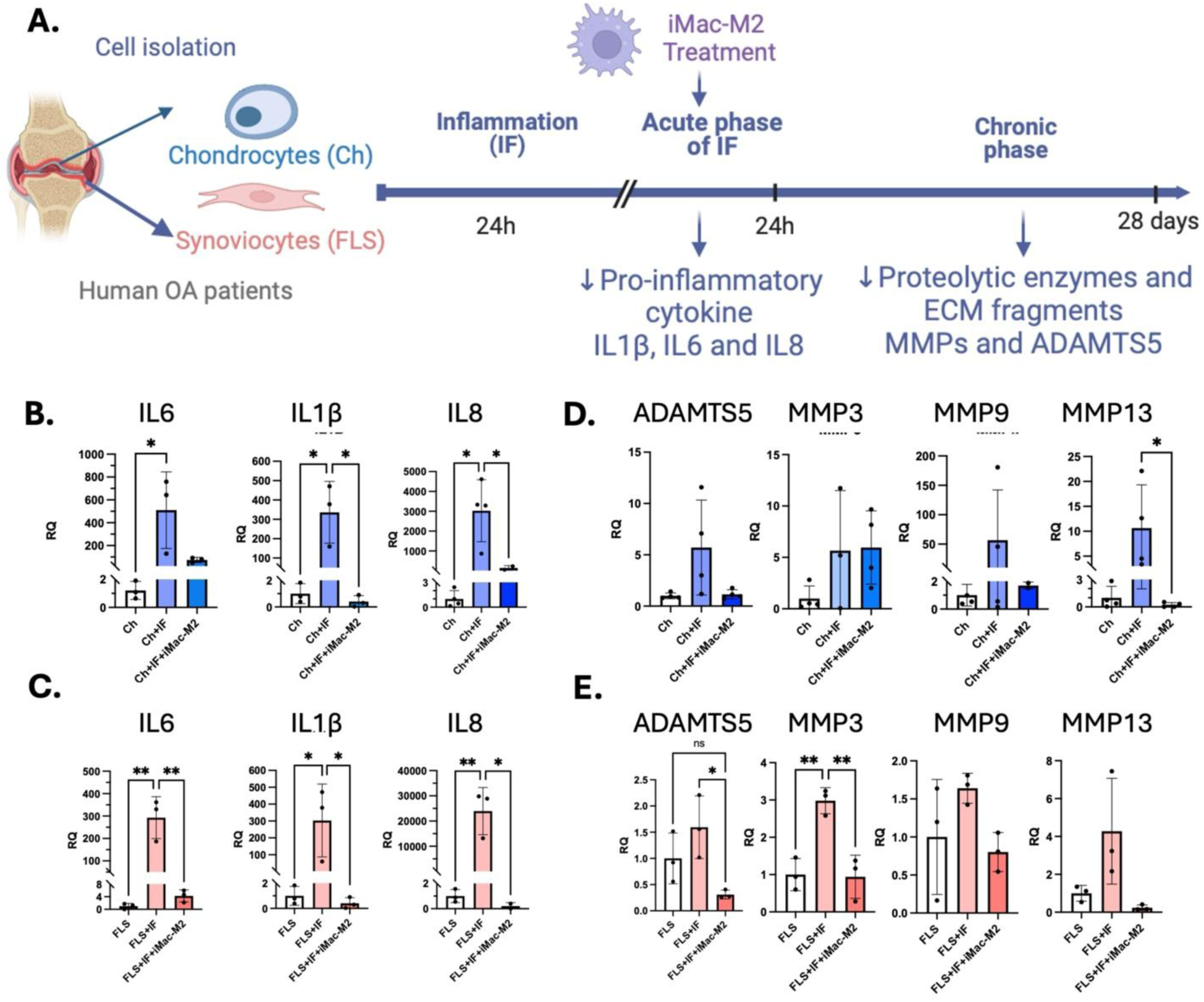
iMac-M2 exert anti-inflammatory effects on osteoarthritic chondrocytes and synoviocytes in vitro. (**A**) Schematic representation of the in vitro osteoarthritis model. Primary chondrocytes (Ch) and fibroblast-like synoviocytes (FLS) isolated from osteoarthritic patients were exposed to inflammatory stimulation, followed by treatment with iPSC-derived M2 macrophages (iMac-M2). Gene expression changes were assessed during the acute inflammatory phase (24 h) and during the chronic phase (up to 28 days). (**B**, **C**) Gene expression analysis of pro-inflammatory cytokines (IL6, IL1β, and IL8) measured 24 hours after induction of inflammation in chondrocytes (blue) and FLS (peach), respectively, showing reduced inflammatory signaling following iMac-M2 treatment. (**D**, **E**) Gene expression analysis of matrix-degrading enzymes (ADAMTS5, MMP3, MMP9, and MMP13) measured 14 days after inflammation in chondrocytes (blue) and FLS (peach), indicating reduced expression of proteolytic enzymes associated with cartilage degradation after iMac-M2 treatment. Data are presented as mean ± SD with individual data points shown. N = 3, N represents the number of independent culture wells from a single co-culture experiment, using primary chondrocytes and fibroblast-like synoviocytes isolated from the same patient donor and iMac-M2 derived from iPSC line CS0007iCTR-n5. Statistical comparisons were performed using one-way ANOVA with Tukey’s multiple-comparisons test. Statistical significance is indicated as *P < 0.05, **P < 0.01; ns, not significant.

To assess longer-term effects, expression of matrix-degrading enzymes was analyzed after prolonged co-culture. iMac-M2 treatment was associated with reduced expression of ADAMTS5, MMP3, MMP9, and MMP13 in chondrocytes (Fig. 4D) and FLS (Fig. 4E).

Collectively, these data show that iMac-M2 treatment attenuates inflammatory cytokine expression and reduces matrix-degrading enzyme expression in human chondrocytes and FLS in vitro.

### Joint response to iMac-M2 treatment in a PTOA rat model

Immunocompetent rats underwent destabilization of the medial meniscus (DMM) surgery to induce PTOA, followed by intra-articular administration of iMac-M2 or PBS control one-week post-surgery (Fig. 5A). Blood samples were collected before injection and one week after treatment to assess systemic IgM levels. Systemic IgM concentrations remained comparable between PBS- and iMac-M2-treated animals before and after injection (Fig. 5B), indicating that iMac-M2 administration did not induce detectable changes in circulating IgM levels during the early post-injection period.

**Fig 5.**
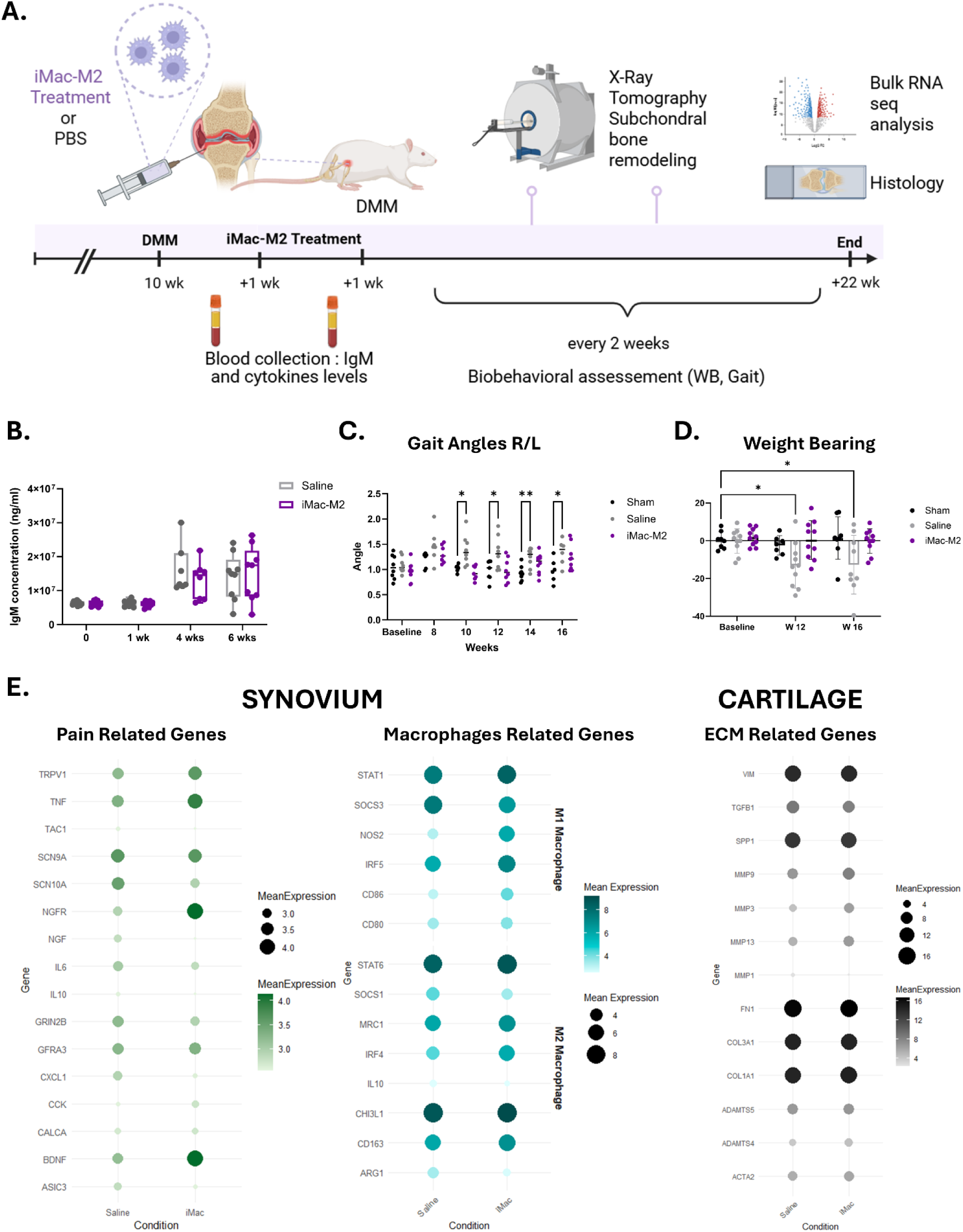
In vivo administration of iMac-M2 in immunocompetent rats following DMM surgery. (**A**) Schematic representation of the experimental design. Destabilization of the medial meniscus (DMM) surgery was performed on the right knee of each rat to induce post-traumatic osteoarthritis. One week after surgery, animals received an intra-articular injection of PBS or iMac-M2 cells. Blood samples were collected before and one week after injection to measure IgM and cytokine levels. Biobehavioral assessment tests (BBT), including acetone, gait analysis, and von Frey testing, were performed every two weeks. X-ray microtomography was used to evaluate subchondral bone remodeling. At the experimental endpoint (16 weeks post-DMM), tissues were collected for histological analysis (OARSI scoring), whole-mount immunolabeling and clearing for nerve and macrophage imaging, and bulk RNA sequencing. (**B**) Systemic IgM levels (ng/mL) measured before injection and one week after treatment with PBS (control) or iMac-M2 (N=8-10). (**C**) Gait analysis showing right and left gait angles measured longitudinally in Sham, Saline, and iMac-M2 groups. (**D**) Weight-bearing analysis evaluating hind limb load distribution between groups over time. In (B-D), n represents the number of animals per group. Data in (C) and (D) are presented as mean ± SD with individual animals shown. Statistical comparisons in (B-D) were performed using repeated-measures two-way ANOVA with Tukey’s multiple-comparisons test. Statistical significance is indicated as *P < 0.05, **P < 0.01. (**E**) Bulk RNA sequencing analysis of synovium and cartilage highlighting gene expression changes associated with pain-related pathways, macrophage-related genes in synovium, and extracellular matrix–related genes in cartilage to assess gene expression, n = 3, n represents the number of animals per group.

Functional outcomes were assessed longitudinally using gait analysis and weight-bearing measurements. PBS-treated animals exhibited a progressive increase in gait asymmetry between the operated and contralateral limbs over time (Fig. 5C). In contrast, animals receiving iMac-M2 displayed reduced gait asymmetry, with values remaining closer to baseline levels throughout the follow-up period. Consistent with these observations, weight-bearing analysis showed that iMac-M2-treated rats maintained a more balanced load distribution between hind limbs compared to PBS-treated controls (Fig. 5D), suggesting improved recovery following treatment. cartilage harvested from PBS- and iMac-M2-treated animals (Fig. 5E). In synovial tissue, expression levels of pain- related genes including TRPV1, TNF, TAC1, and SCN9A were similar between groups, whereas NGFR, NGF, and IL6 showed lower expression in iMac-M2-treated animals, consistent with attenuated inflammatory and nociceptive signaling compared to PBS controls. Genes associated with macrophage-related pathways, such as SOCS3 and MRC1, exhibited comparable expression levels between treatment groups.

In cartilage, expression of extracellular matrix-related genes, including MMP3, MMP9, and MMP13, did not differ significantly between PBS- and iMac-M2-treated animals (Fig. 5E), indicating limited early transcriptional remodeling of cartilage at the analyzed time point.

### iMac-M2 preserve joint structure and limit degeneration in a PTOA model

Longitudinal micro-computed tomography was performed to assess subchondral bone remodeling following DMM surgery in Sham, DMM-Saline, and DMM-iMac-M2 groups at weeks 10 and 22 (Fig. 6A). Three-dimensional reconstructions and coronal cross-sections revealed marked subchondral bone degeneration in saline-treated DMM joints, characterized by trabecular disorganization and bone loss, which progressed between weeks 10 and 22. In contrast, joints treated with iMac-M2 exhibited preservation of subchondral bone architecture over time, with structural features more closely resembling those observed in Sham controls (Fig. 6A).

**Fig 6.**
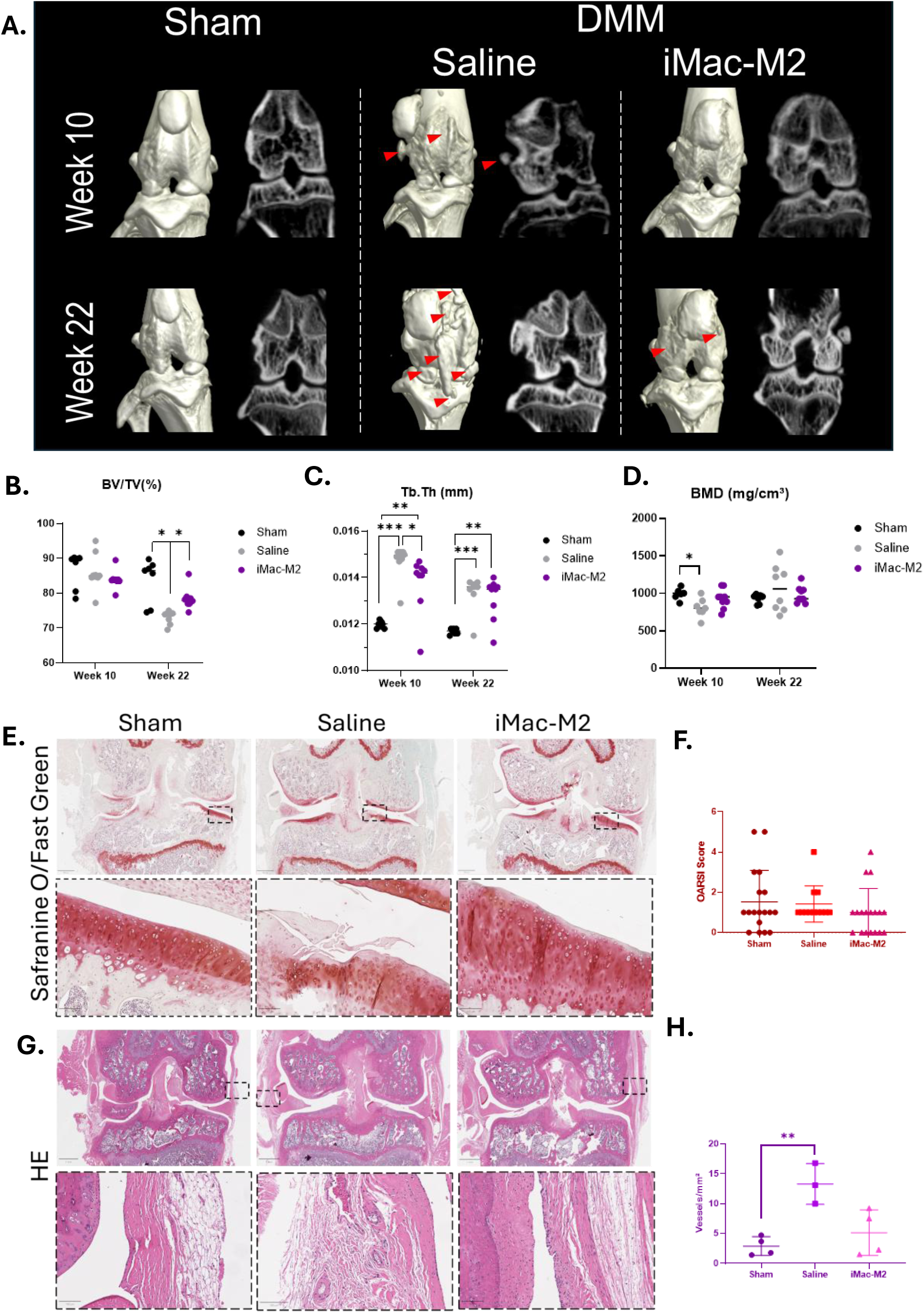
iMac-M2 treatment preserves subchondral bone structure and reduces cartilage degeneration and synovial inflammation in the DMM rat model. (**A**) Longitudinal micro–computed tomography (micro-CT) imaging of subchondral bone structure in Sham, DMM-Saline, and DMM–iMac-M2 groups at weeks 10 and 22 post-surgery. Three-dimensional reconstructions and coronal cross-sections highlight subchondral bone degeneration (red arrows) in the saline-treated DMM group, whereas iMac-M2 treatment preserves bone architecture over time. (**B**) Quantification of bone volume fraction (BV/TV) at weeks 10 and 22. (**C**) Quantification of trabecular thickness (Tb.Th) at weeks 10 and 22. (**D**) Quantification of bone mineral density (BMD) at weeks 10 and 22. (**E**) Histological assessment of cartilage integrity using Safranin O/Fast Green staining in Sham, DMM-Saline, and DMM–iMac-M2 groups. (**F**) Quantification of cartilage degeneration using the OARSI scoring system based on Safranin O/Fast Green–stained sections. (**G**) Hematoxylin and eosin (H&E) staining of synovial tissue illustrating synovial inflammation. (**H**) Quantification of synovial vascular density Data are presented as mean ± SD with individual data points shown. In (B), (C), and (D), N = 8–10, where N represents the number of animals per group. In (F), n = 20– 24, where N represents the number of sections scored, obtained from n = 8-10 animals per group. In (H), n = 3–4, n represents the number of animals per group. Statistical comparisons were performed using two-way ANOVA with Tukey’s multiple-comparisons test in (B–D) and one-way ANOVA with Tukey’s multiple-comparisons test in (F) and (H). Statistical significance is indicated as *P < 0.05, **P < 0.01, ***P < 0.001, ****P < 0.0001.

Quantitative micro-CT analysis supported these observations. Bone volume fraction (BV/TV) was significantly reduced in saline-treated DMM joints compared to Sham at both time points, whereas iMac-M2-treated joints showed higher BV/TV values relative to saline-treated animals at weeks 10 and 22 (Fig. 6B). Similarly, trabecular thickness (Tb.Th) was markedly decreased in the saline group, while iMac-M2 treatment was associated with partial preservation of trabecular thickness at both time points (Fig. 6C). Bone mineral density (BMD) followed a comparable pattern, with reduced values in saline-treated DMM joints and higher BMD measurements in iMac-M2-treated joints, particularly at the early time point (week 10) (Fig. 6D).

Cartilage integrity was subsequently evaluated using Safranin O/Fast Green staining (Fig. 6E). Saline-treated DMM joints exhibited clear signs of cartilage degeneration, including surface irregularities, reduced proteoglycan staining, and structural disorganization. In contrast, cartilage from iMac-M2-treated joints showed better preservation of tissue architecture and proteoglycan content, as indicated by stronger Safranin O staining intensity compared to saline- treated controls. These histological findings were supported by OARSI scoring, which revealed lower cartilage degeneration scores in the iMac-M2 group relative to saline-treated animals (Fig. 6F).

Synovial tissue remodeling was assessed by hematoxylin and eosin staining (Fig. 6G). Saline-treated DMM joints displayed pronounced synovial thickening and hyperplasia, consistent with ongoing synovial inflammation. In comparison, synovial tissue from iMac-M2-treated joints appeared less hypertrophic and exhibited a more organized synovial lining, suggesting reduced inflammatory remodeling. Quantification of vascular density demonstrated a significant increase in synovial neovascularization in saline-treated DMM joints compared to Sham, whereas iMac- M2 treatment was associated with a reduction in vascular density, approaching levels observed in Sham joints (Fig. 6H).

## Discussion

In this study, we show that iMac-M2 can modulate key inflammatory and structural features associated with PTOA. Through a combination of in vitro and in vivo experiments, we demonstrate that iMac-M2 retain a stable M2 phenotype under inflammatory stress, suppress the production of pro-inflammatory cytokines and matrix-degrading enzymes in patient-derived chondrocytes and synoviocytes, and preserve joint structure and function in a rat model of PTOA induced by DMM. Notably, iMac-M2 treatment was associated with behavioral improvement in weight-bearing and gait symmetry, without inducing systemic immunogenicity, supporting the translational potential of this immunomodulatory approach in an immunocompetent setting.

### A robust and reproducible platform for macrophage therapy and iMac-M2 phenotypic stability

The stepwise differentiation protocol yielded iMac-M2 with high purity (Fig. 1), as evidenced by robust expression of CD163 and CD206, key markers of the M2 phenotype, consistent with previously published iPSC-to-macrophage differentiation strategies, including the protocol described by Park and collaborators.^41^ Whereas most previous studies evaluate macrophage polarization over a limited number of differentiation cycles or short culture windows, our study extends this analysis across 24 independent harvests (corresponding to approximately 12 weeks of sustained production), providing a more stringent assessment of long-term phenotypic stability and reproducibility (Fig. 1 C-E). Notably, selected cultures were maintained for up to 44 weeks (Fig. S1), during which a gradual decline in iMonocyte output was observed, while macrophage differentiation and M2 polarization were preserved.

The consistency of this process across multiple iPSC lines highlights the robustness of the platform, a critical requirement for therapeutic development (Fig. S1). Importantly, iMac-M2 retained their anti-inflammatory characteristics even after prolonged culture, suggesting that iMacs may acquire a reinforced polarization state that is less susceptible to environmental reprogramming (Fig. 3C-F). This behavior contrasts with primary human macrophages, which exhibited pronounced plasticity under inflammatory challenge, losing M2 markers and acquiring pro-inflammatory traits in response to TNF-α and LPS (Fig. S2).

Sustained activation of STAT3- and STAT6-associated transcriptional programs may contribute to this phenotypic stability,^42^ as these pathways are known to reinforce M2 identity while suppressing NFκB-driven inflammatory signaling.^43^ Consistent with this interpretation, our gene expression analyses revealed stimulus-dependent regulation of STAT-associated pathways under inflammatory conditions (Fig. 3G-I). This enhanced stability is particularly relevant in the context of chronic joint inflammation, where macrophages are continuously exposed to pro-inflammatory cues.

In line with our findings, several recent studies have reported the generation of iMacs for inflammatory and degenerative indications.^44–46^ However, these studies primarily focus on macrophage generation, short-term phenotypic characterization, or disease models outside the musculoskeletal system. By contrast, our work extends this growing body of literature by demonstrating that iMac-M2 remain phenotypically stable under sustained inflammatory stress (Fig. 3) and exert functional and structural benefits in a clinically relevant PTOA model (Fig. 5-6).

### Resilience of iMac-M2 to inflammatory and pro-inflammatory stimuli

In co-culture experiments, iMac-M2 effectively suppressed inflammatory cytokine production (IL-6, IL-1β, IL-8) in chondrocytes and synoviocytes, while also reducing the expression of matrix-degrading enzymes MMP3, MMP9 and ADAMTS5^47^ (Fig. 4). These findings are consistent with previous reports demonstrating that M2 macrophages promote inflammation resolution and tissue protection through IL-10– and TGF-β–dependent mechanisms.^48^ Importantly, our data extend these observations by showing that iMac-M2 maintain these anti-inflammatory and anti-catabolic effects under sustained inflammatory conditions, including exposure to IL-1β, TNF-α, and LPS (Fig. 3). This is a critical point, as several studies have reported that macrophages, including M2-like populations, may undergo phenotypic drift in inflammatory environments. In contrast to iMac-M2, primary human macrophages exposed to identical stimuli (Fig. S2) in our study exhibited a pronounced loss of M2 markers and increased expression of M1- associated markers, highlighting the greater susceptibility of primary macrophages to inflammatory reprogramming. Our results instead suggest that iMac-M2 maintain their anti-inflammatory phenotype despite sustained exposure to inflammatory cues (Fig. 4C–F). These findings suggest that iMac-M2 possess an intrinsic resistance to pro-inflammatory re-education, a property that may be particularly advantageous for therapeutic applications in PTOA, where inflammation is persistent and multifactorial.

### In vivo efficacy of iMac-M2 in a rat model of PTOA

In the DMM-induced PTOA model, a single intra-articular administration of iMac-M2 one week after surgery resulted in measurable improvements in joint function, as evidenced by improved gait (Fig. 5C), and weight-bearing behavior (Fig. 5D). These functional outcomes are clinically relevant, as altered gait patterns and load redistribution are strongly associated with pain and disease progression in PTOA.^49–51^ At the structural level, iMac-M2 treatment preserved subchondral bone architecture and attenuated synovial inflammation, two early pathological features increasingly recognized as key drivers of PTOA progression (Fig. 6). In contrast, effects on cartilage gene expression were comparatively modest, suggesting that iMac-M2 primarily modulate inflammatory and osteo-synovial components rather than directly inducing cartilage regeneration at the analyzed time points. This observation is consistent with emerging concepts that early inflammatory and subchondral bone changes may precede irreversible cartilage degeneration, reinforcing the importance of early immunomodulatory intervention.^52^

### Implications for clinical translation and future directions

These findings support the potential of iMac-M2 as an immunomodulatory strategy targeting early inflammatory drivers of PTOA. From a translational perspective, future work should focus on defining optimal delivery strategies, including dosing regimens, timing of administration, and the potential benefit of repeated intra-articular injections. In addition, combination approaches integrating iMac-M2 with cartilage-protective or regenerative therapies may further enhance therapeutic efficacy. Manufacturing considerations will also be critical for clinical development. The scalability, batch-to-batch consistency, and compliance with good manufacturing practice (GMP) standards associated with iMac-M2 represent key factors that must be addressed to enable clinical translation.

### Limitations

This study has several limitations. The persistence, biodistribution, and long-term fate of iMac-M2 following intra-articular injection were not directly assessed and will require dedicated tracking approaches. Only a single injection administered at one post-injury time point was evaluated, limiting conclusions regarding optimal therapeutic windows. Moreover, although male rats were selected due to their faster PTOA progression, sex-specific differences in immune responses and disease trajectory warrant future investigation. Finally, validation of these findings in large-animal models will be necessary to better approximate human joint biomechanics and disease progression before clinical application.

## Methods

### 1. iPSC cell culture and differentiation

#### 1.1. iPSC maintenance and expansion

Human iPSC lines were obtained from the Cedars-Sinai Core facility in accordance with institutional IRB approval (No. 42241). The primary line used in this study was a 007iCTR healthy control line (CS0007iCTR-n5) from the Cedars-Sinai Biomanufacturing Center repository, reprogrammed from peripheral blood mononuclear cells of a 60- year-old male donor using non-integrating episomal vectors, with no associated clinical diagnosis or known pathogenic mutations. The line is not deposited in a public registry and is available from the Cedars-Sinai Biomanufacturing Center.

iPSCs were expanded on Matrigel™-coated plates (BD Biosciences) (0.08 mg/well in 6-well plates) and fed every other day with cGMP mTeSR™ Plus medium (StemCell Technologies). Passaging was performed using ReLeSR™ (StemCell Technologies) following the manufacturer’s protocol, and cells were cryopreserved in CryoStor CS10. iPSCs were maintained between passages 26 and 40, and the macrophage differentiation experiments reported in this study were performed using cells at passage 37.

All four lines completed the differentiation protocol and generated iMac-M2 populations (Supplementary Fig. S1). CS0007iCTR-n5 was selected as the primary line for the studies reported here because it produced the highest iMonocyte yields, with greater viability and more abundant monocyte output per harvest, and was therefore used for all subsequent characterization and experiments.

#### 1.2. iPSC identity and quality control

The identity and quality of the iPSC line were confirmed prior to use. Clearance of episomal reprogramming vectors was verified by EBNA genomic DNA PCR. Line identity was authenticated by short tandem repeat (STR) profiling, which showed 100% identity to the parental donor line, and genomic integrity was confirmed by G-banded karyotyping, which revealed a normal 46,XY male karyotype. Cultures tested negative for mycoplasma. Pluripotency was verified by immunocytochemistry for OCT4, SOX2, NANOG, SSEA-4, TRA-1-60, and TRA-1-81 together with alkaline phosphatase staining, and tri-lineage differentiation potential was confirmed by embryoid body formation followed by hPSC Scorecard analysis demonstrating endoderm, ectoderm, and mesoderm induction.

#### 1.3. iPSC to iMacrophage differentiation

iPSC-derived monocytes (iMonocytes) were generated as previously described ^53^. Briefly, maintained iPSCs were differentiated into monocytes through a sequential protocol involving cytokine stimulation. iPSCs were exposed to M-CSF (50 ng/mL, R&D Systems, Minneapolis, MN, USA) and IL-3 (20 ng/mL, R&D Systems) for 7 days in serum-free differentiation medium, with medium changed every 2 days and cells maintained at 37°C in a humidified 5% CO₂ incubator. The resulting iMonocytes were characterized by flow cytometry, showing expression of monocyte markers (CD14, CD11b).

iMonocytes were then polarized into M2 macrophages (iMac-M2) as previously described ^41^. Briefly, 1 × 10⁶ iMonocytes were incubated in M2 polarization medium containing IL-4 (20 ng/mL, R&D Systems), IL-6 (20 ng/mL, R&D Systems), and IL-13 (20 ng/mL, R&D Systems) for 48 hours. The medium was replaced every 2 days, and cells were cultured at 37°C in a humidified 5% CO₂ incubator. The differentiated iMac-M2 were characterized by flow cytometry for macrophage-specific markers (CD206, CD163). After differentiation, iMac-M2 were used for intra-articular injection as part of the in vivo experiment.

### 2. Flow cytometry/functional assays

#### 2.1. Flow cytometry analysis

To assess both monocytes and macrophages levels for induction and polarization efficiency, iPSC-derived cells induced the day of harvest, 5- and 7-days post differentiation were lifted with Enzyme (1X) TrypLE™ Express (5 minutes at 37°C) and washed three times with FACS buffer (2% BSA and 0.1% sodium azide in PBS). Cells were either left unstained or stained with anti-human antibody (Table 1), for 20 minutes at 4°C in the dark. After washing, cells were resuspended in FACS buffer, and data were acquired using a BD LSR Fortessa analyzer (BD Biosciences). Gating strategy included exclusion of debris and doublets, followed by live cell gating and marker-specific thresholds defined using fluorescence-minus-one controls. Analysis was performed using FlowJo software (FlowJo LLC).

**Table 1.**
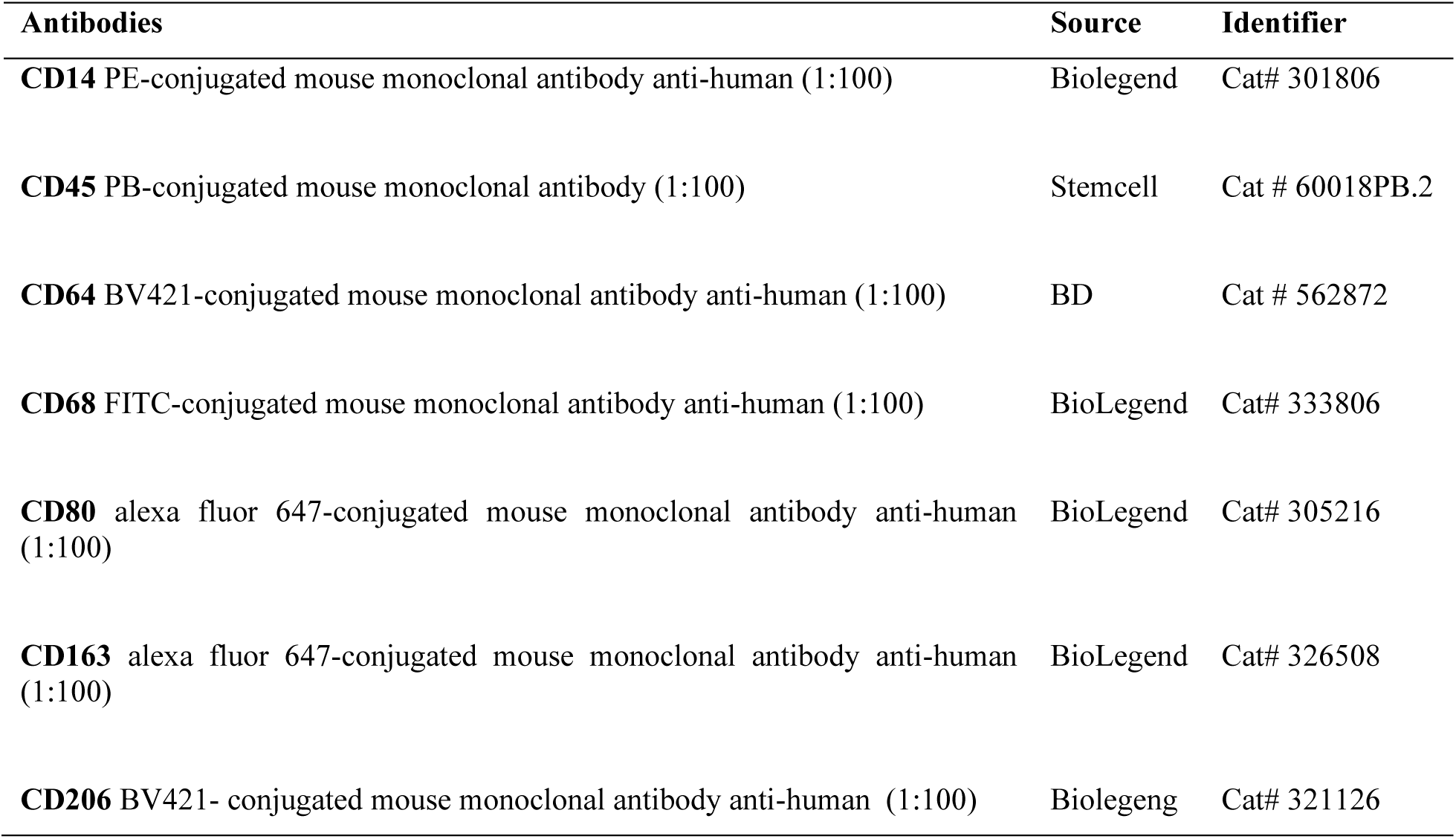
List of antibodies used for Flow cytometry.

#### 2.2. Phagocytosis Assay with Zymosan Particles

The functionality of iMac-M2 cells was assessed by phagocytosis using *Saccharomyces cerevisiae* Zymosan A BioParticles (Alexa Fluor 594 conjugated) (Invitrogen). Zymosan particles were reconstituted in PBS with 2 mM sodium azide and sonicated to disrupt aggregates. Particles were added to adherent iMac-M2 cells at a 2:1 ratio (particles to cells) in serum-free RPMI medium.

Phagocytosis was allowed for 30 minutes at 37°C, followed by washing with PBS, trypan blue to quench surface-bound particles, and another PBS wash. Cells were detached with 0.25% trypsin/0.5 mM EDTA at 4°C for 1 hour, then centrifuged, fixed in 4% formaldehyde, and analyzed by flow cytometry (FACSCanto II, BD Biosciences, San Jose, CA, USA) to assess the phagocytosis. Phagocytosis was quantified as the percentage of Alexa Fluor 594–positive iMac-M2 cells by flow cytometry after removal of non-internalized particles during washing steps and quenching of extracellular fluorescence with trypan blue. Data were processed using FlowJo software, with fluorescence-negative cells used to set a threshold for quantifying the percentage of cells that ingested Zymosan.

#### 2.3. Bulk RNA sequencing

Total RNA was extracted from samples using the RNeasy Plus Kit (Qiagen, Hilden, Germany) according to the manufacturer’s instructions. RNA concentration was measured using the Qubit™ RNA High Sensitivity Assay Kit (Invitrogen, Carlsbad, CA, USA), and RNA integrity was assessed with the Agilent TapeStation system using RNA ScreenTape (Agilent Technologies, Santa Clara, CA, USA).

For library preparation, 1000 ng of total RNA per sample (26 µL input volume) was used. Ribosomal RNA depletion was performed using the RiboCop rRNA Depletion Module (Lexogen, Vienna, Austria), followed by library construction with the xGen™ Broad Range RNA Library Prep Kit (Integrated DNA Technologies, IDT, Coralville, IA, USA). Indexing PCR amplification was carried out using the Normalase™ Unique Dual Indexing Primer Plate (IDT, Coralville, IA, USA) with 10 amplification cycles.

Final libraries were quantified using the Qubit™ dsDNA High Sensitivity Assay Kit (Invitrogen, Carlsbad, CA, USA) and assessed for size distribution and quality using the Agilent TapeStation system with High Sensitivity D1000 ScreenTape (Agilent Technologies, Santa Clara, CA, USA). Sequencing was performed on an Illumina NovaSeq platform (Illumina, San Diego, CA, USA) using a 10B flow cell and a 300-cycle sequencing kit, at a loading concentration of 130 pM, with a 1% PhiX control spike-in.

Raw sequencing reads were processed using a standard RNA sequencing analysis pipeline. Reads were aligned to the rat reference genome (Rnor_6.0), and gene-level read counts were generated. Differential gene expression analysis was performed using the DESeq2 package in R. Adjusted p-values were calculated using the Benjamini–Hochberg method, and genes with an adjusted p-value < 0.05 were considered differentially expressed.

### 3. Coculture of iMac-M2 and Chondrocytes or Synoviocytes (FLS)

#### 3.1. Chondrocytes and FLS isolation

Human cartilage and synovial tissue samples were obtained from 5 Cedars-Sinai patients undergoing joint replacement surgery, following informed consent in accordance with institutional IRB approval (No. 52234). Cartilage tissue was finely minced and digested sequentially with collagenase P 0.1% (w/v) 14h in DMEM/F12 medium at 37°C with agitation to isolate chondrocytes. Synovial tissue was processed separately by digesting with collagenase P 0.1% (w/v) in DMEM/F12 medium under the same conditions to isolate synoviocytes.

After digestion, cells were filtered through a 70µm mesh, washed in PBS, and counted. Chondrocytes and synoviocytes were seeded separately in complete culture media (DMEM/F12 supplemented with 10% FBS, 1% penicillin-streptomycin and 0.1% L-ascorbic) at a density of 1 × 10⁶ cells per 100 mm plate. Cells were cultured for 48 hours to allow adhesion, followed by biweekly media changes to support growth and maintenance.

#### 3.2. Inflammation activation

Inflammatory media was prepared by adding 10ng/mL of IL-1β and 100ng/mL of lipopolysaccharide (LPS) to the chondrocyte and synoviocyte culture media. Chondrocytes and FLS were seeded at a density of 1 × 10⁴ cells per well in 24-well plates and allowed to adhere prior to stimulation. Cells were then exposed to inflammatory media for 24 hours to induce an osteoarthritis-like inflammatory phenotype. After 24 hours of stimulation, inflammatory media was removed and replaced with fresh culture medium prior to co-culture experiments.

To capture both early and sustained inflammatory responses, gene expression analyses were performed at defined time points corresponding to acute and chronic phases of the in vitro osteoarthritis model. The acute inflammatory phase was assessed 24 hours after stimulation, whereas longer-term responses associated with osteoarthritis progression were evaluated at 14 days in chondrocytes and 28 days in FLS.

#### 3.3. Direct Coculture

Following 24 hours of inflammation induction, iMac-M2 cells were directly co-cultured with chondrocytes. iMac-M2 cells were added at a density of 1×10^5^ cells per well to wells containing pre-seeded chondrocytes or FLS.

#### 3.4. FACS – Fluorescence Activated Cell sorting

iMac-M2 cells were stained with a CD14 PE-conjugated mouse monoclonal antibody (anti-human, 1:100 dilution, BioLegend), and then sorted from chondrocytes and synoviocytes at multiple time points (Day 1, Day 3, Day 7, and Day 14) using a BD Influx cell sorter (BD Biosciences) with a 100 µm nozzle, set to the ’1.0-drop purify’ mode. After sorting, the cells were collected in RLT buffer for gene expression analysis.

#### 3.5. Gene expression analysis

Following FACS, cells were collected, and total RNA was extracted using the RNeasy Plus kit (Qiagen). The RNA was then reverse transcribed into cDNA using the High-Capacity cDNA Reverse Transcription Kit (Applied Biosystems). cDNA was amplified using qPCR with TaqMan® gene expression assays. The threshold cycle (Ct) value for 18S rRNA was used as an internal control, employing the TaqMan® FAM/MGB probe system (4333760F, Thermo Fisher). Gene expression levels were calculated using the Livak method, with fold change determined as 2-ΔΔCt, as previously described.

### 4. In vivo injection of iMac-M2

#### 4.1. Destabilization of Medial Meniscus (DMM)

A total of 30 immunocompetent male Sprague-Dawley rats (8 weeks old) underwent DMM surgery under general anesthesia in accordance with institutional IACUC approval (No. 010218). Briefly, the procedure involved making a medial parapatellar incision on the right knee to access the joint capsule. Using a microsurgical scalpel, the medial menisco-tibial ligament was transected to destabilize the medial meniscus. The contralateral left knee was left unoperated to avoid systemic confounders. Following the procedure, the joint capsule and skin were sutured, and the animals were closely monitored during recovery. Post-operative care included administration of analgesics and ensuring appropriate wound healing to minimize discomfort.

#### 4.2. Injection of iMac-M2

One week after DMM surgery, animals were assigned to receive an intra-articular injection of either iMac-M2 or vehicle control. iMac-M2 cells were harvested, washed twice in sterile phosphate-buffered saline (PBS), and resuspended at the desired concentration immediately prior to injection.

Rats were anesthetized using isoflurane, and the skin over the knee joint was disinfected. Intra-articular injections were performed under aseptic conditions using a 30-gauge needle inserted through the patellar tendon into the joint space. A total volume of 25 µL containing 1 × 10⁶ iMac-M2 cells was injected into the operated knee. Sham-operated animals underwent the same surgical exposure without ligament transection and received intra-articular injection of sterile PBS under identical conditions.

Animals were randomly assigned to treatment groups, and behavioral and histological analyses were performed by investigators blinded to treatment.

After injection, animals were allowed to recover under close monitoring and were returned to their cages once fully awake. No adverse events related to the injection procedure were observed. Animals were subsequently followed longitudinally for behavioral, imaging, histological, and molecular analyses as described below.

### 5. Serum analysis: IgM ELISA

To assess systemic inflammation markers, blood samples were repetitively collected from each animal at baseline (day of surgery), and at 1-, 2-, 8-, and 16-weeks post-surgery via tail vein puncture. A 0.8 mL serum separator tube (Greiner Bio One MiniCollect, Monroe, NC, USA) was used for collection. The samples were then centrifuged at 2000×g for 10 minutes. Finally, the serum was separated and stored at -80°C until analysis. Serum IgM levels were measured using an ELISA kit (MBS2510638; MyBioSource, San Diego, CA, USA) to evaluate the immune response to iMac- M2 administration. Standard curves were performed in accordance with the standard values indicated by the manufacturer of the ELISA kits. All data sets were exported to Excel, and concentrations were calculated using interpolation in Prism 8 software (GraphPad, La Jolla, CA, USA).

### 6. Analysis of PTOA Development

#### 6.1. Behavioral assessment of locomotor function and weight bearing

Gait analysis was performed to assess functional impairment and disease progression following DMM surgery. Rats were trained to walk along a straight corridor lined with filter paper prior to testing. At baseline and at 48 hours, 2-, 8-, and 16-weeks post-surgery, non-toxic red paint was applied to the forelimbs and blue paint to the hind paws, and footprint patterns were collected. Footprints were analyzed to quantify hindlimb gait parameters, including stride length, inter-step (sway width) distance, and hind paw angle. These measurements were used to assess gait symmetry and limb use between the injured and contralateral hind limbs over time.

Weight-bearing distribution was assessed to evaluate joint discomfort and functional loading of the hind limbs. Rats were placed in a transparent enclosure and allowed to move freely without restraint. Hindlimb weight distribution was recorded using the Dynamic Weight Bearing Test 2.0 (Bioseb, France) which is a pressure-sensitive force platform system equipped with high-sensitivity sensors and synchronized overhead video monitoring. Weight distribution between the injured and contralateral hind limbs was continuously recorded over a 5-minute period and used as an index of joint incapacityand pain-related behavior.

#### 6.2. X-ray microtomography analysis of subchondral bone

In-vivo micro computed tomography (μCT) scans were used to assess the bone mineral density (BMD) and structural parameters of the knee joints. Scans were performed using a Quantum GX2 microCT scanner (PerkinElmer, Waltham, MA, USA) at a voltage of 55kVp and a current of 145mA, with an integration time of 200ms and a voxel size of 35μm. The entire knee joint was scanned to capture structural details of the bone. Imaging analyses were performed using eXplore MicroView software (General Electric Healthcare, Milwaukee, WI) to contour the region of interest and quantify key architectural parameters including bone volume, total volume, bone volume fraction (BV/TV), and mean density. These parameters were used to evaluate the effects of iMac-M2 administration on the progression of post-traumatic osteoarthritis (PTOA) in the DMM rat model. Image reconstruction and quantitative analyses were performed using MicroView software (Parallax Innovations Inc., Ilderton, ON, Canada).

#### 6.3. Histology

Following sacrifice, the knee joints were harvested, fixed in 4% formaldehyde, dehydrated through graded ethanol, and embedded in paraffin. Five-micron-thick sections were cut from the paraffin blocks.

Sections were stained with hematoxylin and eosin (H&E) to assess overall tissue morphology and synovial architecture. Safranin O/Fast Green staining was performed to evaluate cartilage integrity and glycosaminoglycan (GAG) content.

Synovial vascularization was quantified on H&E-stained sections. Vascular density was assessed by manually counting blood vessels within the synovial tissue using QuPath software (version 0.5.0, University of Edinburgh, Edinburgh, UK). Quantification was performed on standardized regions of interest from multiple sections per joint by investigators blinded to treatment, and vascular density was expressed as the number of vessels per mm².

## Statistical Analysis

Statistical analyses were completed using the GraphPad Prism 10 (GraphPad Software, San Diego, CA, USA) and R software. Data are presented as mean ± standard deviation (SD) unless otherwise stated. Comparisons between two groups were performed using unpaired two-tailed student’s t-tests. Comparisons involving multiple groups or time points were analyzed using one-way or two-way ANOVA with post-hoc multiple-comparison tests. For longitudinal behavioral assessments, repeated-measures ANOVA was used. The value of P < 0.05 was considered statistically significant. Statistical significance was defined as * P <0.05, ** P < 0.01, *** P <0.001, and **** P < 0.0001. For bulk RNA sequencing analyses, differential gene expressions were analyzed using DESeq2 with Benjamini-Hochberg correction for multiple testing, and adjusted P values < 0.05 were considered significant.

## Acknowledgements

The authors would like to acknowledge the Cedars-Sinai Biobehavioral Core, Cedars-Sinai biobank, and the Cedars-Sinai Imaging Core for their support in this work. This study was supported by internal funding sources.

