## Supplemental Figures for "Modulating Inflammation in Post-Traumatic Osteoarthritis using iPSC-derived Anti-inflammatory Macrophages"

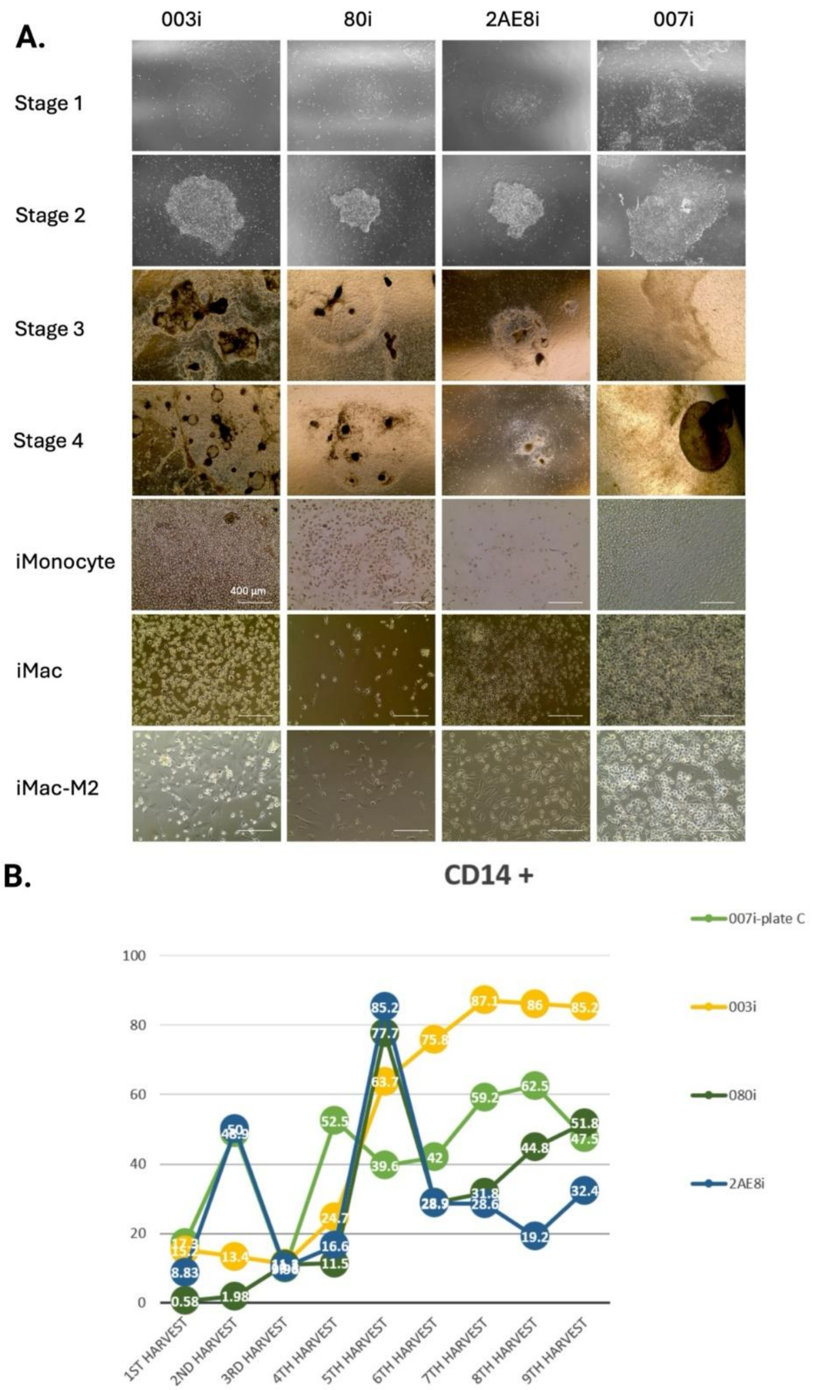


***Fig. S1. Differentiation of multiple iPSC lines into iMac-M2 macrophages demonstrates the robustness of the differentiation protocol.*** *(A) Representative brightfield images showing the stepwise differentiation of four independent iPSC lines through successive stages of the protocol, including early differentiation stages (stages 1–4), iMonocyte production, macrophage differentiation (Mac), and polarization into anti-inflammatory macrophages (iMac-M2). (B) FACS data showing percentage of CD14 + cells across harvest timepoints in four different iPSC cell lines (n = 4). N represents the number of independent iPSC lines; each line was taken through a differentiation experiment. Each point represents a single harvest from one line.*

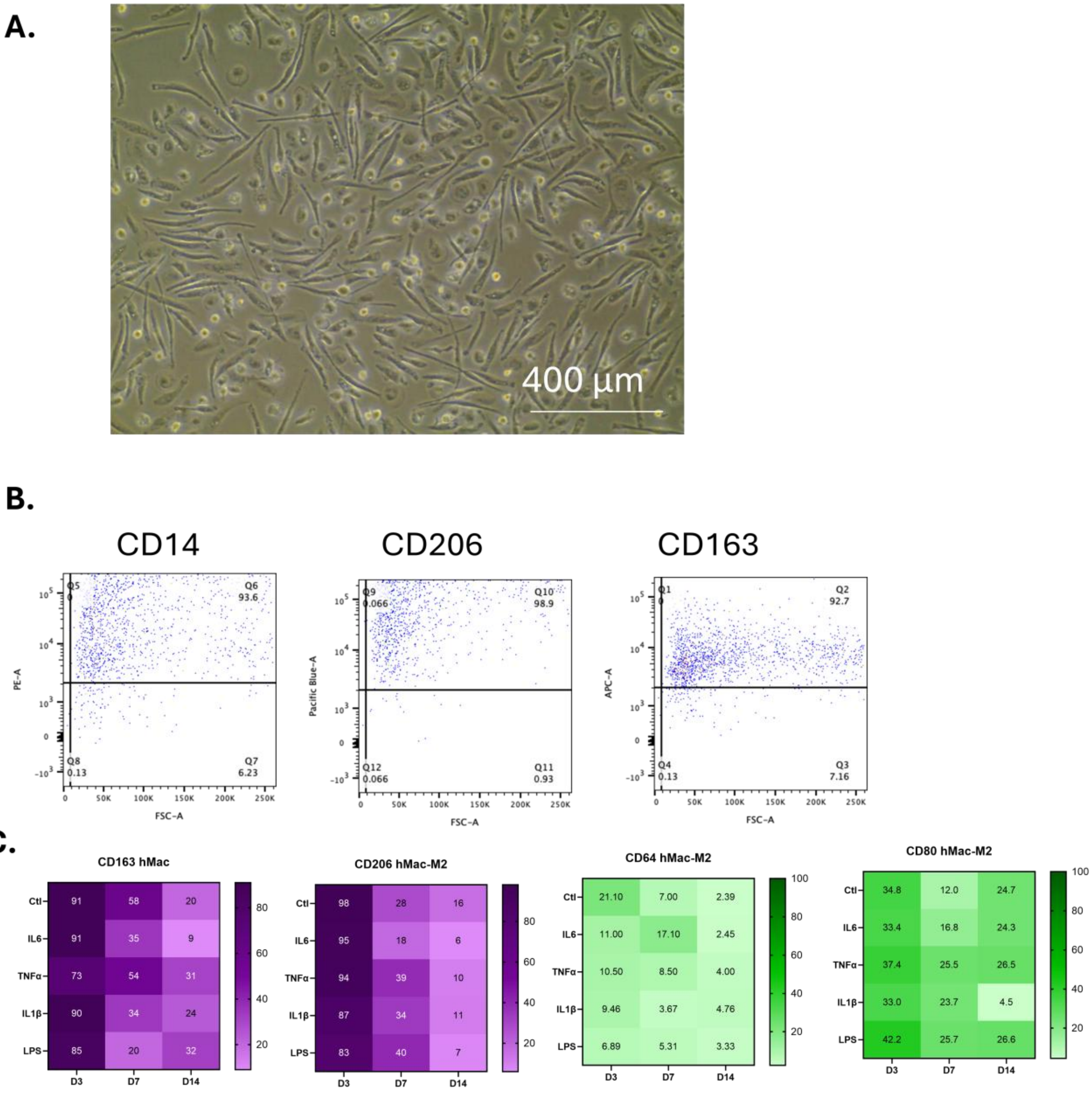


*__Fig. S2. Characterization and inflammatory response of purified human macrophages.__ (A) Representative brightfield image showing the morphology of purified human macrophages (hMacrophages) following M2 polarization (scale bar, 400 µm). (B) Flow cytometry plots illustrating the expression of macrophage and M2-associated markers CD14, CD206, and CD163 after polarization. (C) Heatmap analysis showing the expression levels of M2 markers (CD163, CD206) and M1-associated markers (CD64, CD80) following exposure to inflammatory stimuli (IL6, TNFα, IL1β, or LPS) at day 3, day 7, and day 14, indicating a reduction in M2 marker expression under inflammatory conditions.*

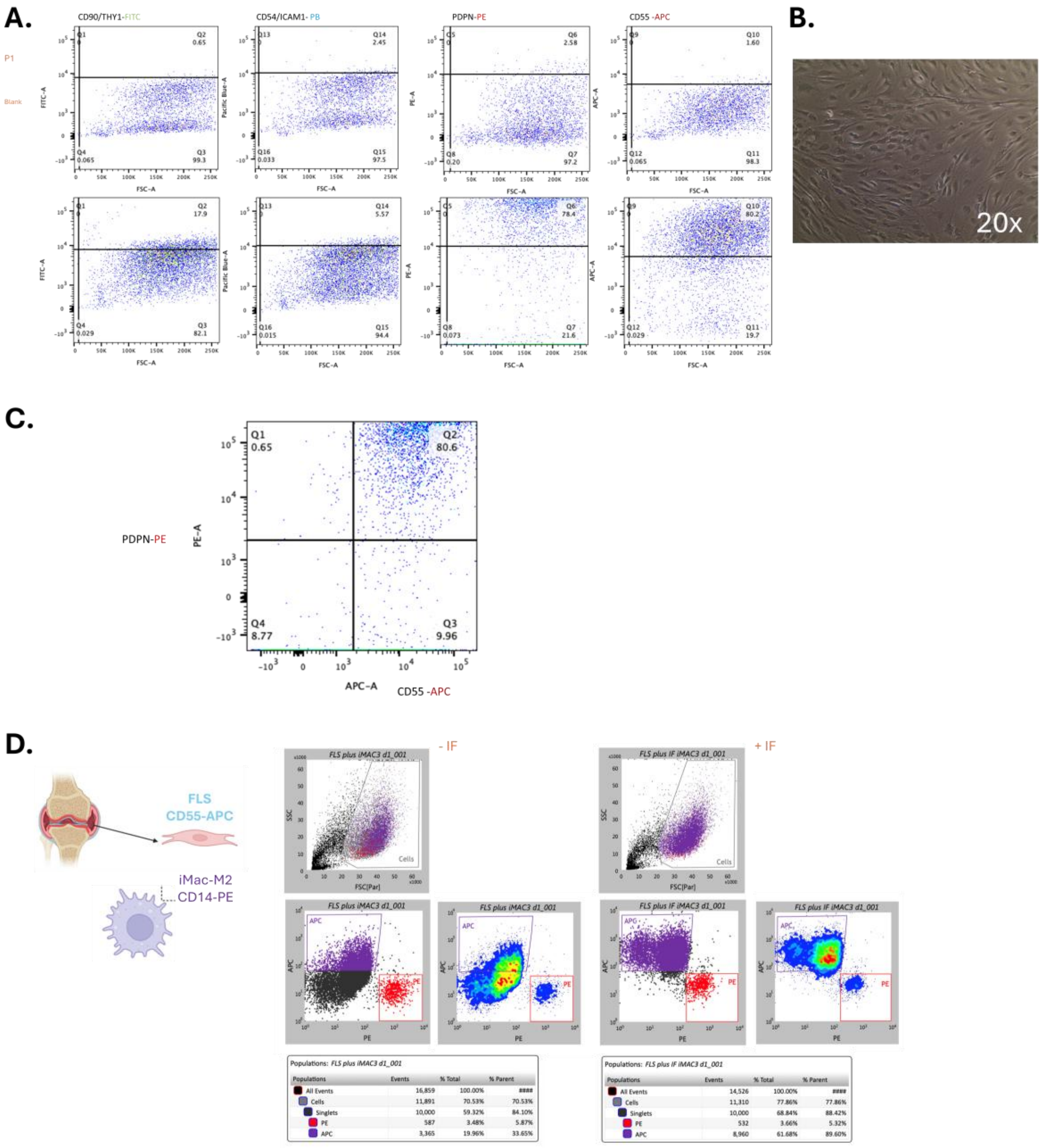


***Fig. S3. Flow cytometric characterization and sorting of fibroblast-like synoviocytes (FLS) and iMac-M2 in co-culture experiments.*** *(**A**) Representative flow cytometry plots showing the expression of fibroblast-associated markers CD90 (THY1), CD54 (ICAM-1), PDPN, and CD55 in fibroblast-like synoviocytes (FLS) isolated from osteoarthritic synovium (patient 088). Brightfield microscopy (20×) illustrates the characteristic spindle-shaped morphology of cultured FLS. (**B**) Flow cytometry analysis demonstrating a PDPN$^{+}$CD55$^{+}$ double-positive phenotype, confirming the fibroblast-like identity of synoviocytes. (**C**) In co-culture experiments, FLS (CD55$^{+}$) and iMac-M2 (CD14$^{+}$) were distinguished by flow cytometry under both non-inflammatory (–IF) and inflammatory (+IF) conditions. Prior to gene expression analysis, cells were sorted using fluorescence-activated cell sorting (FACS) based on these markers to selectively isolate FLS and exclude iMac-M2 cells. This strategy ensured that subsequent transcriptomic analyses reflected cell-specific responses of synoviocytes or chondrocytes without contamination from macrophage-derived RNA.*
